# Environmental microbes promote phenotypic plasticity in *Drosophila* reproduction and sleep behavior

**DOI:** 10.1101/2022.11.26.518045

**Authors:** Mélisandre A. Téfit, Tifanny Budiman, Adrianna Dupriest, Joanne Y. Yew

## Abstract

The microbiome has been hypothesized as a driving force of phenotypic variation in host organisms that is capable of extending metabolic processes, altering development, and in some cases, conferring novel functions that are critical for survival (1-5). Only a few studies have directly shown a causal role for the environmental microbiome in altering host phenotypic features. To directly assess the extent to which environmental microbes induce variation in host life history traits and behavior, we inoculated axenic *Drosophila* with microbes isolated from two different field sites and generated two populations with distinct bacterial and fungal profiles. We show that microbes isolated from environmental sites with modest abiotic differences induce large variation in host reproduction, fatty acid levels, stress tolerance, and sleep behavior. Importantly, clearing microbes from each experimental population removed the phenotypic differences. The results support the causal role of environmental microbes as drivers of host phenotypic variation and potentially, rapid adaptation and evolution.

## Introduction

Most animals acquire a cohort of commensal microbes from environmental sources, the composition of which is shaped in part by abiotic variables including temperature, humidity, and pH (6-9). As such, host organisms of the same species can harbor highly divergent microbial communities depending on the local environment conditions. For example, American pika populations that reside at different elevations exhibit microbiomes with significantly different levels of taxonomic diversity (10). One fascinating consequence of microbiome variation is that it increases the potential for phenotypic variation, at times resulting in new physiological functions for the host (11, 12). This phenomenon has been observed in bean bugs that gain resistance to pesticides by incorporating a pesticide-inactivating bacterium, *Burkholderia*, from the local soil environment (3). Similarly, woodrats exposed to toxic creosote plants are able to metabolize harmful phenols due to an enrichment of environmentally-acquired *Actinobacteria* in their gut microbiome (4). Notably, this extended metabolic phenotype is not observed in individuals of the same species that live in a habitat devoid of creosote plants. In both cases, the uptake of specific microbes from the local environment allowed hosts to adapt and thrive in response to local stressors by conferring novel physiological functions.

Considering the myriad roles that the microbiome plays on the gut-brain axis in animals (13-16), microbial communities acquired from different environments could potentially alter various physiological and behavioral features of the same host species in different ways. To directly test this possibility, we inoculated axenic *Drosophila melanogaster* with microbes isolated from one of two field sites with modest differences in elevation, rainfall, and temperature. Each experimental population exhibited different reproductive abilities, stress resistance, and lipid profiles. Unexpectedly, inoculation with different microbial communities also led to distinct patterns of diurnal sleep behavior. The differences in microbial profile and the attendant phenotypic contributions persisted through multiple generations in a laboratory setting. In addition, the observed effects were completely reversible and the phenotypic differences between the two populations were eliminated once microbes were cleared from the host.

## Methods

### Laboratory diet

Flies were raised on a minimal yeast-sucrose diet (named “YS-”) comprised of 1% agar, 2.5% sucrose, 8% organic cornmeal, and 1.5% inactive yeast (w/v). No antibacterial or antifungal agents were added to the food in order to retain as much of the complexity in wild-isolated microbiome as possible.

### Generation of axenic flies

Axenic flies were generated from an isogenic *D. melanogaster* CantonS line (BL9517) according to established protocols (17). Briefly, flies were allowed to deposit eggs onto fruit juice agar for 12-16 hrs. The eggs were washed with 5% hypochlorite with 0.05% Tween-20 with gentle agitation. After 5 min, the bleach solution was removed and the eggs washed twice with phosphate buffered saline with filter sterilized 0.05% Tween-20 (PBST). The eggs were dispensed into sterile food with antibiotics (50 μg ampicillin, 50 μg kanamycin, 50 μg tetracycline, and 15 μg erythromycin per mL of media) and raised in an incubator at 25 °C with 50% humidity under a 12: 12 hr day/ night cycle. To test for the absence of microbes, three flies from each vial were individually homogenized in PBS and the homogenate was plated on both MRS and YPD media. No detectable growth was observed on either media after 72 hrs.

### Wild microbiome collection

We collected *Drosophila spp*. from two sites (designated “W1” and “W2”) within the Waimea Valley located on the north shore of Oʻahu, Hawaiʻi (**Fig. 1**). The location contains multiple climate and habitat types, thus providing a wide array of environmental conditions and site-specific microbiome compositions. A microbiome survey of aquatic, terrestrial, and organismal features of the site was recently described (18). To generate the inoculant, newly collected wild male flies were transferred to YS-food vials and raised at 25 °C with 50% humidity under a 12: 12 hr day/ night cycle. After two days, the flies were removed and the top layer of the seeded food scraped and homogenized in 1 mL of sterile PBS. The resulting slurry was used to inoculate fresh YS-food (50 μL per vial, three replicate vials per site). Five to seven axenic *D. melanogaster* flies (2 males and 3 – 5 females) were transferred to inoculated vials. To slow the growth of mold, flies were raised at 19 °C, 50% humidity, 12:12 hr day/ night cycle. The W1 and W2 populations were flipped onto fresh YS-food every 3 weeks in a biosafety cabinet.

**Figure 1.**
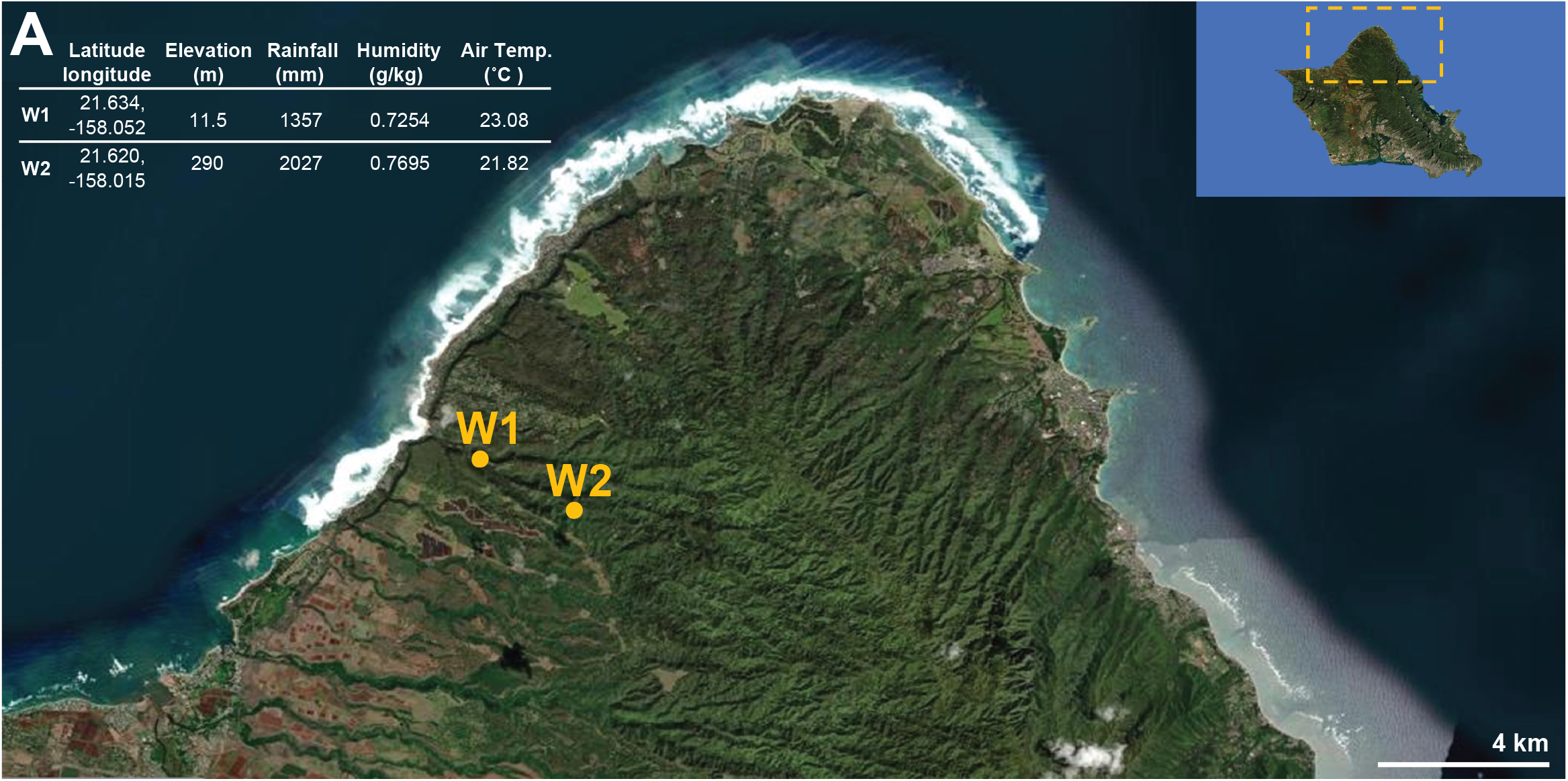
Drosophilids were collected from the Waimea Valley located on the North Shore of Oʻahu. Average annual measures of environmental conditions are provided for the two sample sites, W1 and W2. Figures in modified form and data were sourced from Climate of Hawaiʻi website (72-74).

### High throughput sequencing

Fly surfaces were sterilized with 2 washes in 95% EtOH followed by 2 washes in sterile water. Individual flies were placed in tubes containing 1.4 mm ceramic beads (Qiagen; MD, USA) and ATL buffer from PowerMag Bead Solution (Qiagen) and homogenized using a bead mill homogenizer (Bead Ruptor Elite, Omni, Inc; GA, USA) and extended vortexing for 45 min at 4 °C. Following an overnight proteinase K treatment (2 mg/ mL) at 56 °C, DNA was extracted according to manufacturer’s instructions. Bacterial diversity was characterized using by PCR amplification of the 16S rRNA gene with primers to the V3-V4 region (515F: GTGYCAGCMGCCGCGGTAA; 806R: GGACTACNVGGGTWTCTAAT) (19). Fungal diversity was characterized using primers to the internal transcribed spacer (ITS1f: CTTGGTCATTTAGAGGAAGTAA; ITS2: GCTGCGTTCTTCATCGATGC) (16). The primers contain a 12 base pair Golay-indexed code for demultiplexing.

PCR reactions were performed with the KAPA3G Plant kit (Sigma Aldrich, MO, USA) under the following conditions: 95 °C for 3 min, followed by 35 cycles of 95 °C for 20 seconds, 50 °C for 15 seconds, 72 °C for 30 seconds, and a final extension for 72 °C for 3 min. The PCR products were cleaned and normalized using the Just-a-plate kit (Charm Biotech, MO, USA). High throughput sequencing (HTS) was performed with Illumina MiSeq and 250 bp paired-end kits (Illumina, Inc., CA, USA). The sequences were demultiplexed into FASTQ files on the Illumina BaseSpace Sequence and processed using the MetaFlow|mics analysis pipeline (20). Read filtering, denoising, and merging were performed by DADA2 using the following parameters: reads shorter than 20 bp and samples with fewer than 5,000 reads were discarded. Paired reads are merged if the overlap is at least 20 bp with a maximum 1 bp mismatch (21). The contigs generated by DADA2 were subsequently processed using MOTHUR (22) and aligned and annotated using the SILVA v138 database (23), FungiDB (24), and MycoBank (25). Chimeric contigs were removed with VSEARCH (26). For both 16S and ITS libraries, operational taxonomic units (OTUs) were clustered at similarity levels of 97% and above. The OTUs with a total abundance below 2 were discarded. Sample normalization was performed by subsampling each sample to the same level (10^th^ percentile of the sample sizes). Finally, the LULU algorithm (27) with default settings was used to remove spurious clusters arising from PCR and sequencing artefacts or intra-individual variability.

### Microbiome data analysis

Community data analysis was performed using MicrobiomeAnalyst (28, 29). To remove low quality features, a low count filter (fewer than 4 counts for at least 20% of its values) and a low variance filter (variance lower than 10%) were applied. We applied a centered log ratio transformation to raw count data. Alpha-diversity (intra population diversity) was evaluated at the 97% OTU level using the Shannon index (which accounts for taxa richness as well as evenness) and Chao1 index (which measures richness only in terms of presence or absence). Beta-diversity (diversity between populations) was evaluated using the Bray-Curtis dissimilarity, a quantitative measure that accounts for taxa abundance, and projected in nonmetric multidimensional scaling (NMDS). Permutational analysis of variance (PERMANOVA) was used to estimate the statistical significance of the differences in community composition. For relative abundance profiling, taxa with fewer than 10 counts were merged. Bacterial and fungal OTUs were classified to genus or the lowest possible taxonomic level.

### Physiological assays

Physiological assays were performed using flies that spanned 3 – 5 different generations for each population (2 bottles were raised per generation for each population).

#### Egg laying

One day old males and females (2-3 flies of each sex) were placed in a fresh food vial and flipped into a new vial every 2 days for a total of 16 days (8 vials). Eggs were scored under 10X magnification. Dark pupae from egg-laying vials were counted every day until 20 days post-pairing. Emerging adults were counted every 2 days for 8 days.

#### Analysis of ovarian tissues

Ovaries from 4-5 d old mated females were dissected in PBS. Flies from 5 different bottles were used for each population. Ovarioles and mature egg chambers (Stages 13-14) were scored under 8X magnification.

#### Mitochondria membrane potential measurement

Ovaries from 3 - 4 d old females were dissected in PBS, incubated in MitoTracker Green FM (200 nM in PBS; Thermo Fisher Scientific Inc., Waltham, MA) for 30 min. After 15 min, tetramethyl rhodamine ethyl ester (TMRE; Sigma Aldrich) was added to the same well (final concentration: 50 nM). Ovaries were washed 3 times in PBS and individual ovarioles were separated, placed on a slide with PBS, and imaged immediately using a confocal laser scanning microscope equipped with a white light laser (TCS Leica SP8 X; Leica Microsystems Inc., Deerfield, IL). The MitoTracker dye was excited at 490 nm and TMRE excited at 552 nm. For each ovariole, 3 – 7 Z-sections, each 1.04 μm thickness, were acquired in counting mode under identical gain, laser power, and threshold settings and using Leica LAS X software (version 3.5.5.19976). The ratio between the pixel intensities of the TMRE and MitoTracker signals was determined by applying the “ratio” operation to regions of interest drawn around each ovariole. Three consecutive Z sections were analyzed and the average ratio calculated for each ovariole.

#### Desiccation resistance

Eight to ten flies (age 3-4 d) from each population were placed in empty food vials (25 × 95 mm). The vials were capped with a foam plug and 3 g of Drierite desiccant (OH, USA) and sealed with parafilm. Ten to twelve replicate vials per population were set up in parallel. Vials were monitored every 30 min for attrition.

### Lipid analysis

Fatty acid extraction: Five male or female flies (3 – 4 d old) of the same sex were placed in a screw-cap vial and the tissue was homogenized in water with ceramic beads using a Bead Ruptor homogenizer at 6 m/sec for 30 sec. Twenty μL of the crude extract was removed for protein quantification by a Bradford protein assay according to manufacturer’s instructions (Thermo Scientific Inc.). Next, chloroform: MeOH (2:1, v/v) spiked with pentadecanoic acid (15 μg/ mL) was added to the extract and the samples was vortexed at 1600 RPM for 3 hrs at 4 °C. The samples were briefly centrifuged for 2 min at 7.5k RCF at 4 °C for 2 min and the lower phase was collected and transferred to a clean vial. Extraction with chloroform was repeated twice more, the fractions pooled, and the solvent evaporated to dryness under N2. For fatty acid esterification, 0.5 N methanolic HCl (Sigma Aldrich, MO, USA) was added to the vial and incubated at 65 °C for 90 min. The solvent was evaporated and samples reconstituted in hexane prior to GCMS analysis. Five replicates were prepared per population.

Cuticular hydrocarbon extraction: For each replicate, five male or female flies (3 - 4 d old) were extracted with hexane (Sigma Aldrich) spiked with 10 μg/ mL of hexacosane (Sigma Aldrich). Five replicates each from the W1 and W2 populations were prepared in parallel.

Gas chromatography mass spectrometry (GCMS): GCMS analysis was performed on a 7820A GC system equipped with a 5975 Mass Selective Detector (Agilent Technologies, Inc., Santa Clara, CA, USA) and a HP-5ms column ((5%-Phenyl)-methylpolysiloxane, 30 m length, 250 μm ID, 0.25 μm film thickness; Agilent Technologies, Inc.). Electron ionization (EI) energy was set at 70 eV. One microliter of the sample was injected in splitless mode and analyzed with helium flow at 1 mL/ min. For fatty acids, the oven was initially set at 50 °C for 2 min, increased to 90 °C at a rate of 20 °C/min and held at 90 °C for 1 min, increased to 280 °C at a rate of 5 °C/min and held at 280 °C for 2 min. For CHC analysis, the oven was initially set at 40 °C for 3 min, increased to 200 °C at a rate of 35 °C/min, increased to 280 °C at a rate of 20 °C/ min, and held at 280 °C for 15 min. The MS was set to detect from m/z 33 to 500. Data were analyzed using MSD ChemStation (Agilent Technologies, Inc.). CHCs were identified on the basis of retention time and electron ionization fragmentation pattern. The abundance of each compound was quantified by normalizing the area under each CHC peak to the area of the hexacosane signal using homebuilt peak selection software (personal correspondence, Dr. Scott Pletcher, Univ. of Michigan). To calculate total CHC levels, individual peak areas were normalized to hexacosane and summed.

### Sleep and activity analysis

Individual 5 – 7 d old male or female flies were placed in a sterile 65 mm glass tube that was capped with sterile fly food on one end. The experiments were performed in a 12:12 hr light-dark cycle for 6 days in an incubator held at 25 °C and 50% humidity. Fly activity was quantified by using the Drosophila Activity Monitoring System (DAMS, Trikinetics, MA, USA) to count the number of photobeam crossings in 1 min bins (30). Sixteen flies from each population were measured in parallel for each experiment. Three independent experiments were conducted using male flies and 2 independent experiments were conducted using female flies. In total, flies from 3 different bottles were used for W1/ W2 experiments. For axenic lines, 2 independent experiments were performed. Activity analysis was performed using PHASE (31) to calculate sleep duration per day, number of sleep bouts per day, average sleep bout, and the number of activity counts per waking minute. Activity data were binned into 30 min intervals. The plots show the 24 hr average of 3 days of measurements (Days 2, 3, 4). Sleep events were defined as periods of inactivity lasting at least 5 consecutive minutes. Flies that did not move for 24 consecutive hours were considered dead and removed from the data set.

### Statistical analysis

Data were analyzed using Prism (v9; GraphPad Software, San Diego, CA).

## Results

### The wild-derived microbiomes of lab flies maintain compositional differences across multiple generations

Lab-raised *Drosophila* were inoculated using microbes associated with drosophilids captured from two different sites in the Waimea Valley, thus generating two populations with distinct microbiomes. High throughput amplicon sequencing analysis revealed significant differences in the bacterial components of W1 and W2 microbial communities in both early (G1-4) and late (G12) generations, both in terms of taxonomic diversity and composition (early: PERMANOVA R^2^=0.25, p-value <0.001, mean stress = 0.117; late: p<0.001, R^2^: 0.346, mean stress = 0.07; **Fig. 2 A-C, Supp. Fig. 1A**). Bacteria from the *Acetobacter, Lactobacillus*, and *Providencia* genera were more abundant in the early microbiome of W1 hosts compared to W2 flies whilst *Gluconobacter* and unclassified genera from the *Acetobacteraceae* family were more abundant in W2 flies. In late generations, the relative abundances of *Providencia* and *Sphingobacterium* differentiated between the two populations (**Supp. Fig. 2**). The bacterial symbiont *Wolbachia* was not detected in early or late generations.

**Figure 2.**
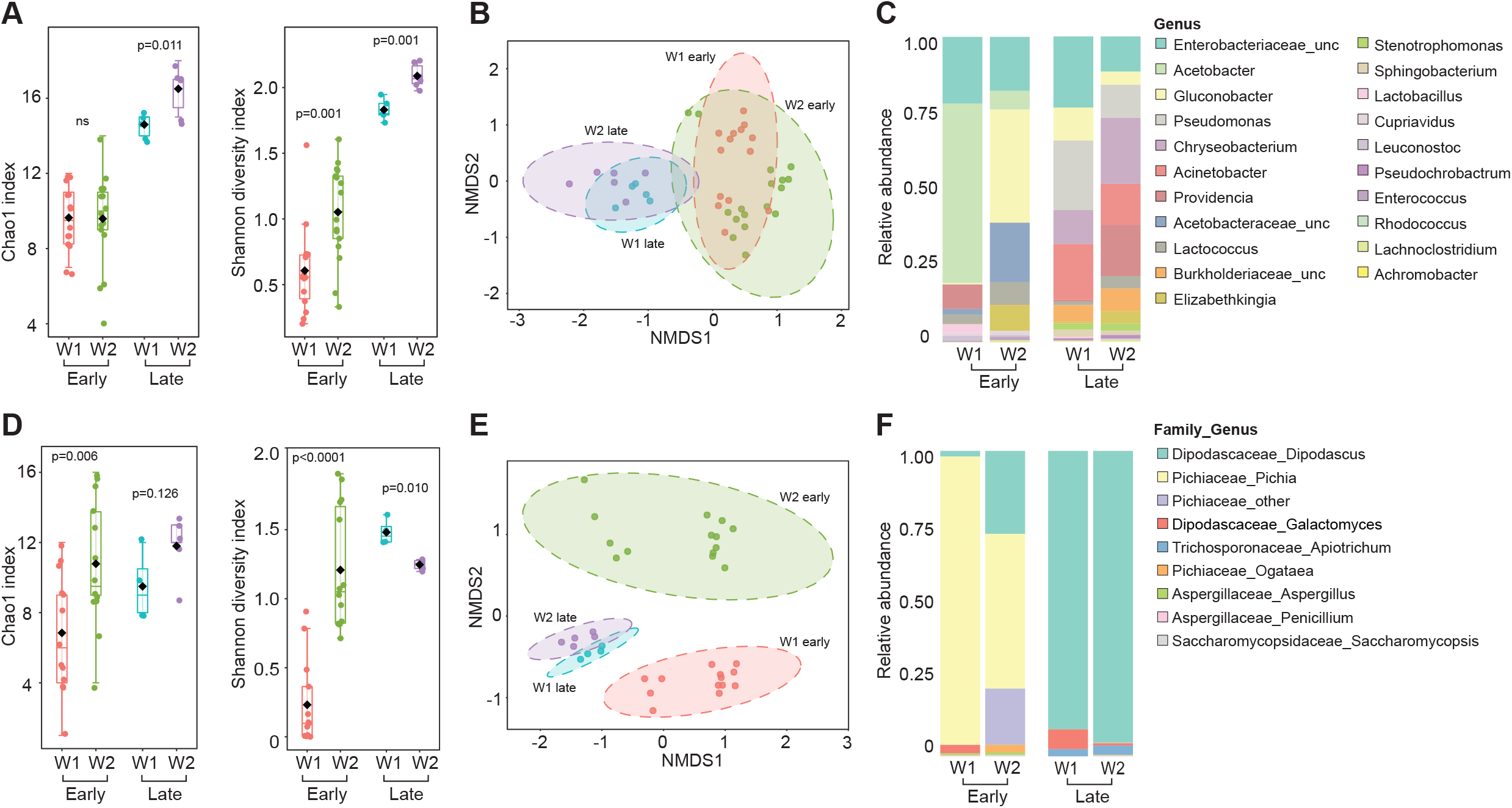
Microbiome community analysis of W1 and W2 populations. (A-C): **(A)** Alpha-diversity analyses of bacterial communities in early (G1-4) and late (G12) W1 and W2 generations. Differences in taxonomic diversity and evenness are maintained under lab rearing conditions. Diversity indices compared using Student’s t-test; ns: not significant. **(B)** Non-multidimensional scaling (NMDS; based on Bray-Curtis distances) reveal distinct bacterial communities in W1 and W2 populations. Ellipses represent significance at 0.05 confidence interval; PERMANOVA: p=0.001; R^2^ = 0.439; mean stress = 0.129. **(C)** Relative abundance plots based on genera. Each column represents the cumulation of 14-17 or 5-6 individual samples for early and late generations, respectively; unc: unclassified genera from the *Enterobacteriaceae, Acetobacteraceae*, and *Burkholderiaceae* families. **(D)** Alpha-diversity analyses of fungal communities in early (G1-4) and late (G12) W1 and W2 generations. Differences in taxonomic diversity and evenness are maintained under lab rearing conditions. Diversity indices compared using Student’s t-test; ns: not significant. **(E)** The NMDS plot of W1 and W2 fungal communities (based on Bray-Curtis distances) reveal different fungal communities in W1 and W2 populations. Ellipses represent significance at 0.05 confidence interval; PERMANOVA: p=0.001; R^2^ = 0.703; mean stress = 0.076. **(F)** Relative abundance plots of 16S and ITS genera. Each column represents the average of 12-14 individuals for early generations and or 4-6 individuals for late generations.

The fungal communities of W1 and W2 populations both in early and late generations were also significantly different in terms of taxonomic diversity and composition (early: PERMANOVA R^2^=0.59, p-value <0.001, mean stress = 0.0702; late: p=0.006; R^2^=0.75; mean stress = 0.05) (**Fig. 2 D-F, Supp. Fig. 1)**. Members of the *Pichiaceae* family, which are common yeast symbionts of drosophilds (32, 33), represented the majority of the fungal community in early generations of both populations. Both *Pichia* and *Galactomyces* genera were more abundant in early generations of W1 flies compared to W2 (**Supp. Fig. 2**). In late generations, the fungal composition of both populations was dominated by *Dipodascus* yeast. There were no significant differences in the relative abundances of individual genera.

### Microbial composition imparts variation in host physiological features

Microbes play a significant role in modulating the gut-brain axis and regulating host metabolism, reproduction, and stress resistance (5, 13, 16, 34-39). To test whether environmental microbes can impart significant variation in physiological features and life history traits, we measured fecundity, fertility, stress resistance, and fat levels in lab populations inoculated with distinct wild microbiomes.

#### Reproduction and development

Flies inoculated with W2 microbes exhibited greater fecundity and faster reproductive rate compared to W1-inoculated flies. The ovaries of W2 flies contained more mature eggs and ovarioles (**Fig. 3A, B**). Although the time to late pupal development and adult emergence was not significantly different between the two populations, W2 flies laid more eggs and had more progeny compared to W1 flies (**Fig. 3D, E, F**). The difference in egg number is due mostly to the accelerated egg laying of W2 flies in the first 11 days after males and females were initially paired. To test the causal role of microbes on reproduction, we cleared W1 and W2 flies of bacteria and fungi by bleaching the eggs from each population and raising flies under sterile conditions. Following microbe removal, the number of ovarioles and mature eggs were no longer different between W1 and W2 populations (**Fig. 3A, C**).

**Figure 3.**
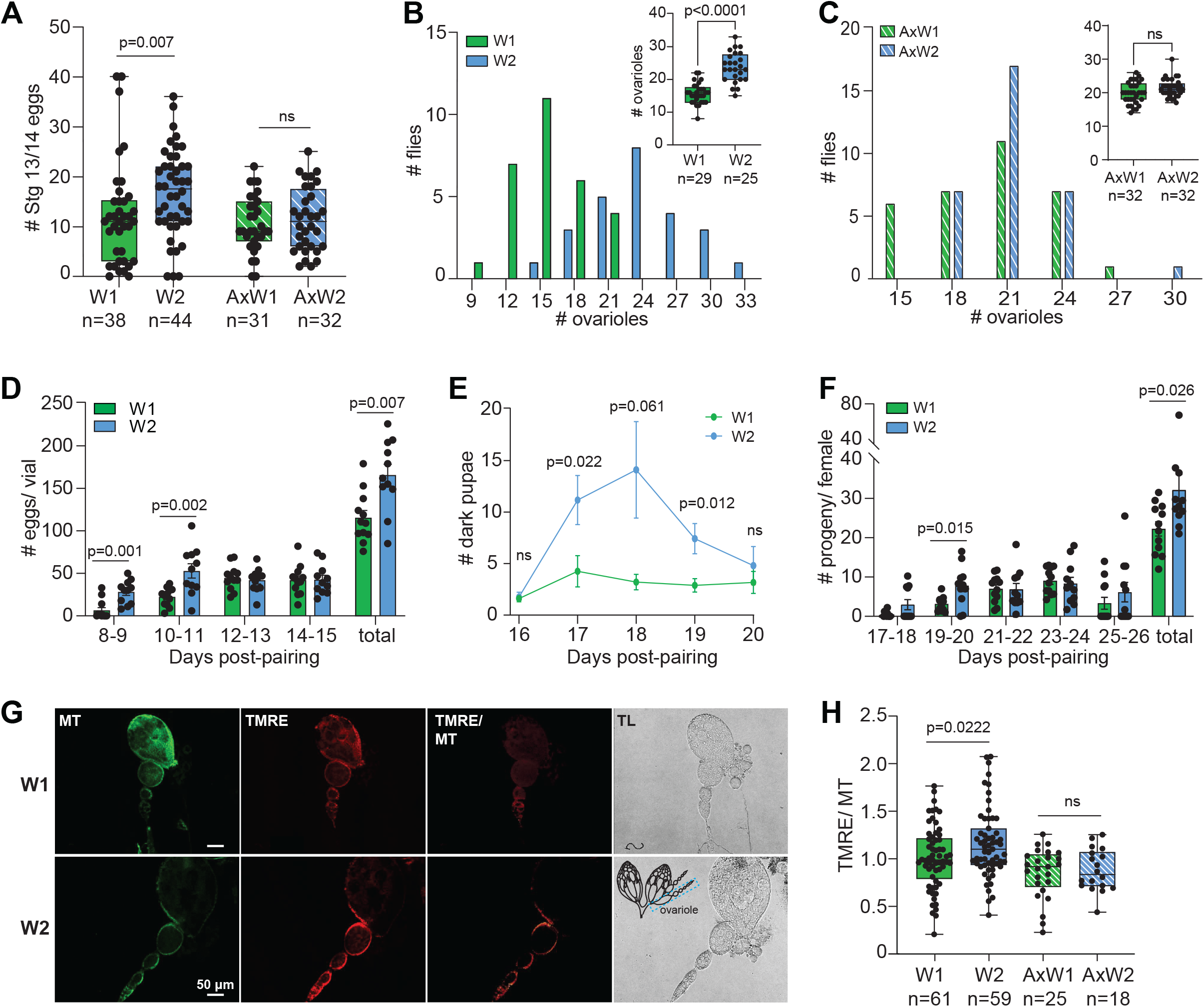
Fertility and fecundity differ between W1 and W2 populations. **(A)** Ovaries from virgin females in the W2 population contain more late stage embryos compared to W1 flies (Mann-Whitney test). Axenic flies generated from both populations (AxW1, AxW2) had similar number of mature eggs. Each point represents an individual fly. Box plots indicate maximum and minimum values; middle bar indicates median. **(B)** Histogram showing the range of ovariole number in W1 and W2 populations. Inset: W2 females have significantly more ovarioles (Mann-Whitney test). **(C)** Histogram showing the range of ovariole numbers in axenic flies. Inset: Axenic W1 and W2 flies have similar number of ovarioles (Mann-Whitney test). **(D)** The W2 females lay more eggs and at a faster rate compared to W1 females (Mann-Whitney). Each point represents the number of eggs laid in a single vial over 48 hrs. Eggs first appeared 8 days after males and females were paired. Bars represent mean ± standard error of the mean (SEM). **(E)** The times to pupation and adult emergence are similar between W1 and W2 populations. There were significantly more dark pupae in W2 vials on Days 17-19 after pairing of males and females and in total (Mann-Whitney test). Each point represents the average of 10-12 vials ± SEM. **(F)** More adults emerged from W2 parents compared to W1 (Mann-Whitney test). Each point represents counts of adults that emerged from a single vial over 48 hrs. Bars represent mean ± SEM. **(G)** Representative confocal images of ovarioles from W1 and W2 populations stained with MitoTracker (MT) and TMRE, an indicator of mitochondrial activity and cellular health; TL: transmitted light. The TMRE/ MT panel shows pseudocolored fluorescence ratio images of TMRE signal intensity (Ex 552 nm) intensity divided by MT signal intensity (Ex 490 nm). The TMRE/ MT ratio of W2 ovarioles is higher than that of W1 ovarioles indicating greater mitochondrial activity in W2 tissues; inset: illustration of an ovary with a single ovariole outlined in blue. **(H)** Quantification of the TMRE/ MT ratio indicates higher mitochondrial potential in W2 ovarioles compared to W1 (Student’s t-test). Ovarioles from corresponding axenic populations show no difference in potential. Box plots indicate maximum and minimum values of square root transformed values; middle bar indicates median.

Because ovarian mitochondrial activity is directly related to oocyte maturation and oogenesis (40-42), we measured mitochondria potential in ovarioles from W1 and W2 flies by using the membrane potential dye TMRE normalized to a marker of mitochondrial mass, MitoTracker. Overall, mitochondrial membrane potential from W2 ovarioles was higher compared to W1 flies (**Fig. 3G, H**). No difference in potential was observed once microbes were removed from the flies (**Fig. 3H**). The higher mitochondrial activity is consistent with our observation that W2 flies are more fecund and fertile than W1 flies.

#### Stress resistance and lipid profiles

Next, we assessed whether microbiome composition can impart differences in desiccation resistance. The W1 flies exhibited consistently higher resistance to desiccation stress compared to W2 flies, for both males and females (**Fig. 4A, B**). Because resistance to environmental stress is correlated with higher fat content (39, 43-45), we compared lipid levels and weight of flies in the two populations. Interestingly, W1 females, but not males, had higher fatty acid levels and weighed more compared to W2 females (**Fig. 4D-G**). We also analyzed the fatty acid composition of these flies but found no significant difference in terms of the relative abundances of mono- and unsaturated fatty acids (data not shown).

**Figure 4.**
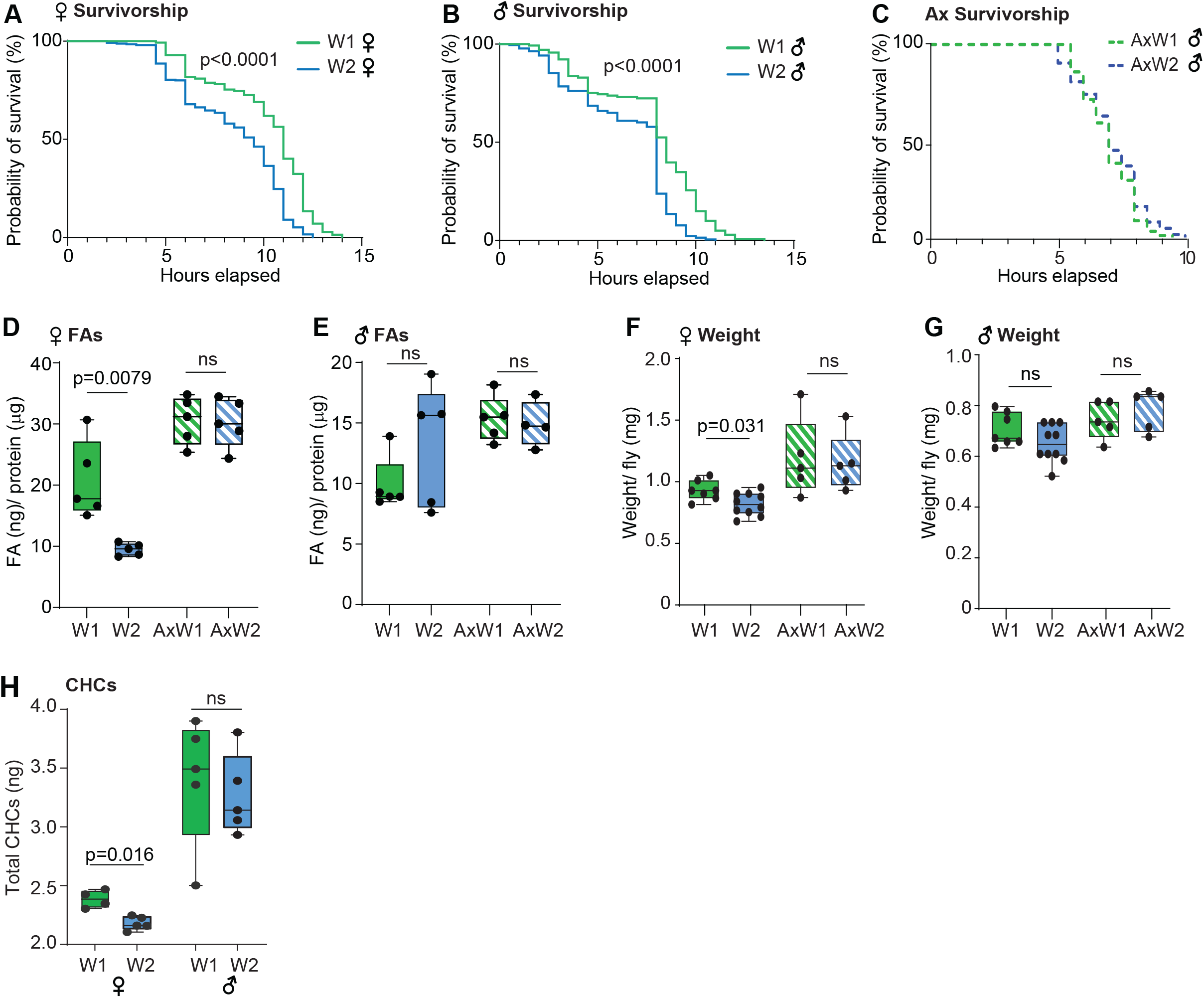
Stress resistance, fatty acid levels, and weight differ between W1 and W2 populations. **(A, B)** W1 females and males survive longer than W2 flies under desiccation stress (Mantel-Cox test; n=16-17 vials for W1; n=23-26 vials for W2). The W1 females have a median survival time of 11 hours compared with 9.5 hours for W2 females. For males, the W1 population has a median survival time of 8.5 hours and 8 hours for W2. **(C)** Axenic W1 (AxW1) and W2 (AxW2) males exhibit similar resistance to stress, both having a median survival time of 7 hours (Mantel-Cox test; n=11 vials for W1; n=13 vials for W2). **(D, E)** Total fatty acid levels (FA) are higher in W1 females compared to W2 females (Mann-Whitney test). Axenic females from each population have similar FA levels. No significant differences in FA levels are found for W1 and W2 males or their axenic counterparts. **(F, G)** The W1 females are heavier than W2 females. Males from W1 and W2 populations exhibit no significant differences in weight. Axenic females or males from each population have similar weights. **(H)** The total abundance of cuticular hydrocarbons (CHC) from W1 females is higher compared to W2 females. No significant differences are found for W1 and W2 males (Mann-Whitney test).

The cuticular layer of insects can also play a significant role in desiccation resistance. The layer of cuticular hydrocarbons (CHCs) that coat the external cuticles of many insects helps to regulate water evaporation, provides protection from infections, and can function as sex or species recognition pheromones (46). Symbiotic microbes contribute to the formation of the cuticular hydrocarbon layer in several species of insects (47-51). We measured cuticular hydrocarbon levels and found that as with fatty acids, W1 females but not males displayed significantly higher levels of CHCs compared to W2 flies. There was no difference in terms of individual CHC types for either population (data not shown). Axenic W1 and W2 flies exhibited similar desiccation resistance and lipid levels, indicating that the microbial composition played a significant role in contributing to the observed differences in stress resistance (**Fig. 4C-G**).

#### Sleep behavior

Sleep physiology, both in health and disease, has been linked to the microbiome and the activity of the gut-brain axis (52, 53). To determine if different microbial communities can shape the structure of sleep-wake cycles, we measured sleep and activity levels in W1 and W2 flies. Surprisingly, our findings show that both male and female W1 flies sleep significantly less during the day and are more active compared to their W2 counterparts (**Fig. 5A-D; Supp. Fig. 3**). This phenotypic change was restricted to daytime sleep and no difference was observed in night time sleep behavior. In addition, diurnal sleep was more fragmented for both W1 sexes, as indicated by a shorter sleep bout duration compared to W2 flies (**Supp. Fig. 3**). Although the W1 females had fewer daytime sleep bouts compared to W2 females, no difference was found between W1 and W2 males (**Supp. Fig. 3**). Importantly, clearing W1 and W2 flies of microbes eliminated the differences in daytime sleep and activity levels (**Fig. 5E-H; Supp. Fig. 3**). We conclude from these findings that microbial composition is the causal factor for variation between the W1 and W2 sleep patterns and that the observed microbe-induced behavioral changes are reversible.

**Figure 5.**
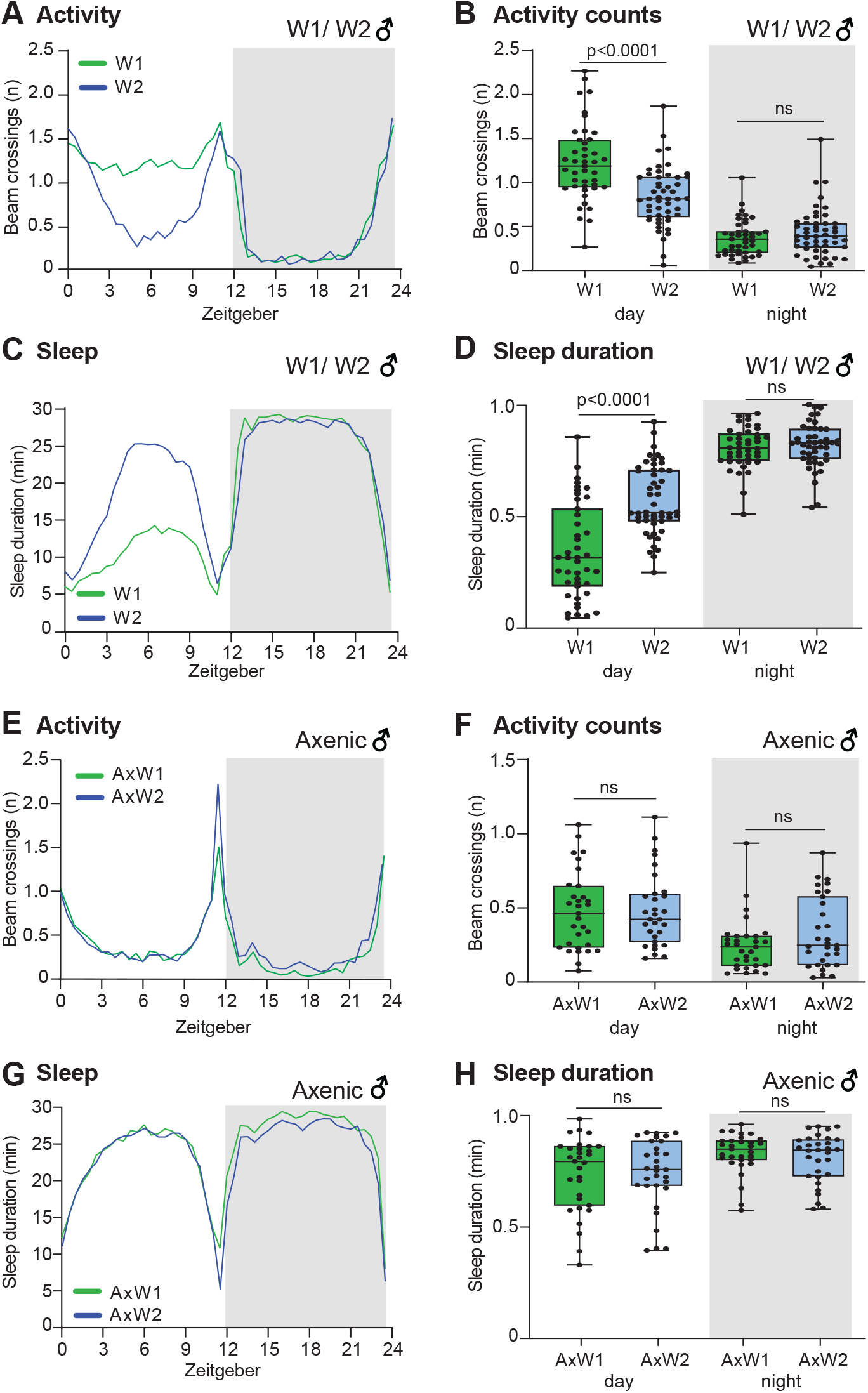
Activity and sleep behavior for W1 and W2 males and axenic populations. **(A)** Daily locomotor activity plots reveal that W1 males are more active during the day compared to W2 males (Mann-Whitney test; n=43 for W1 and n=50 for W2). The two populations exhibit no differences in nighttime activity. The plots show activity counts in 30 min bins averaged over 3 days and summarized in 24 hrs. The gray shading indicates dark periods. **(B)** The average activity count for W1 flies is higher during the day compared to W2 flies but not at night (Mann-Whitney test). Box plots indicate maximum and minimum values; middle line indicates median. Each point represents the average activity count per 30 min bin for a single fly over 3 days. **(C)** The W1 males sleep less during the day compared to W2 males (Mann-Whitney test). No differences are found in nighttime sleep levels. Plots show the average sleep time for W1 and W2 males over 3 days, summarized in 24 hrs. **(D)** The average diurnal sleep duration of W1 flies is significantly shorter than that of W2 males (Mann-Whitney test). Each point represents the sleep duration per 30 min bin for a single fly averaged over 3 days. **(E)** Axenic W1 and W2 males have similar activity profiles during the day and night (Mann-Whitney test; n=43 for W1 and n=50 for W2). **(F)** Axenic W1 and W2 flies have similar diurnal and nocturnal activity counts (Mann-Whitney test). **(G)** Axenic W1 and W2 males exhibit similar sleep profiles during the day and night (Mann-Whitney test). **(H)** Axenic W1and W2 males show no difference in average sleep duration (Mann-Whitney test).

## Discussion

Models of adaptation and phenotypic diversification have traditionally focused primarily on genetic and post-transcriptional mechanisms. However, an increasing number of studies show that the microbiome may also drive evolution drive by altering the genomic architecture of host organisms (54), modulating key physiological traits such as stress tolerance (5, 55), or imparting novel metabolic functions including toxin resistance (4). Our study reveals that microbial communities isolated from sites that exhibit modest differences in elevation (∼280 m) and temperature (1.2 °C) can significantly and distinctly impact critical life history features including reproduction as well as multiple functions modulated by the gut-brain axis, such as stress resistance, sleep, and activity levels. Importantly, once flies were cleared of microbes, the phenotypic differences between the W1 and W2 populations were no longer detected, demonstrating the causal role of the microbiome. The differences in physiological traits and microbiome composition between the experimental populations were sustained over multiple generations of lab rearing despite the absence of an external “wild” microbiome reservoir from which gut microbes can be replenished. We note that unlike the composition of the bacterial community, yeast communities became more similar in later generations. This observation indicates that bacteria may play a greater role in tuning host physiological phenotypes and imparting differences between W1 and W2 populations. Selective elimination of one kingdom using antibiotics or antifungal agents will address this possibility.

Not all of the measured features changed in response to wild microbe inoculation; development time (time to dark pupae formation and adult emergence) was similar between W1 and W2 flies. In addition, whilst microbiome composition largely affected both males and females in similar fashion, only females, but not males, exhibited differences in weight and fatty acid levels. For females of each population, changes in stress tolerance were accompanied by a trade-off in fecundity and fertility: W1 females produced fewer progeny but had greater stress resistance whereas W2 females had a higher reproductive rate but significantly poorer performance under stress. The findings are consistent with well-established patterns of energetic trade-offs between reproduction and survival (44, 56, 57). Notably, lipid metabolism plays a pivotal role in maintaining the balance between the two life history traits: an investment of resources in reproduction results in a depletion of fat stores and shortened lifespan; by contrast, reduced reproduction is associated with increased lipid levels and improved survivorship and stress resilience (39, 43-45). The higher fatty acid and cuticular lipid levels exhibited by W1 females are consistent with the population’s increased stress resistance and lowered reproductive success compared to W2 females. Considering the pivotal role microbes can play in fat storage and nutritional availability (15, 16, 34-37, 58-60), different microbiome communities are likely to influence nutrient levels and modulate the gut-brain axis, resulting in the differential release of neuroactive molecules and hormones that control appetite, the mobilization of energy stores, and reproduction (36, 61, 62). In summary, our observations support a mechanism underlying microbe-induced phenotypic plasticity whereby environmental microbes shift the balance between survivorship and reproduction by influencing lipid metabolism via the gut-brain.

### Sleep and microbiome composition

Amongst the different phenotypes measured, the impact of the microbiome on sleep and activity levels was the most robust and surprising. In the natural world, light and temperature are two well-known environmental cues that strongly regulate sleep-wake cycles (63). For example, daytime siestas are prolonged in natural populations residing in tropical climates and thought to be an adaptation that minimizes exposure to heat (64). Here we show that the microbiome is another environmental factor that contributes to sleep plasticity. Previous studies examining the effect of the gut microbiome on *Drosophila* sleep and activity found no differences when comparing lab-raised male flies to their axenic counterparts (17, 65) and a slight yet inconsistent difference in female conventional vs axenic flies (38). By contrast, our findings reveal that differences in microbiome composition rather than its absence or presence can have a significant effect on daytime sleep. Interestingly, night time sleep was not affected, an observation that may be explained by the fact that the two types of sleep are considered to be distinct in nature, differing in terms of duration and arousal threshold (66). Potential molecular components in *Drosophila* that may link microbiome function to factors that influence day time sleep include the hormone signals ecdysone (67), sex peptide (68), and astrocyte-derived factors including the glucose and lactate transporter *rumple (69)*, the monoamine acetyltransferase *AANAT1 (70)*, and a taurine transporter (71). Each of these signals respond to dietary changes and nutrition levels; as such, it may be the case that different microbiome compositions alter sleep behavior by modulating nutrient availability and, in turn, the release of hormones or metabolites. A metabolome comparison of W1 and W2 populations will help to narrow down the possible mechanisms that underlie the different sleep patterns and possibly identify new pathways that shape sleep behavior.

## Conclusion

Our findings reveal that the environmental microbiome is capable of inducing significant variation in traits that are critical for host fitness and reproduction and potentially, adaptive plasticity. The higher desiccation tolerance exhibited by W1 flies may be an advantage for the drier, warmer conditions of the lower garden site whereas the faster reproductive rate of W2 flies may be adaptive for an environment with limited nutritional resources. The adaptive advantages of the microbe-induced phenotypes remain untested. Moreover, microbial community dynamics under laboratory conditions are likely to be more stable than in natural populations due to limitations placed on host range, diet, and access to different microbial reservoirs. It could be the case that the microbiomes of natural populations fluctuate too much to exert long-term changes in physiology or behavior. Comparing the responses of experimental populations in natural conditions will be a crucial next step for assessing the extent to which environmental microbes contribute to host plasticity and rapid adaptation.

## Acknowledgements

MAT: conceptualization, methodology, investigation, formal analysis, and editing

TB: methodology, investigation, formal analysis, and editing

AD: methodology, investigation, formal analysis, and editing

JYY: conceptualization, methodology, investigation, validation, formal analysis, resources, writing, and editing.

This work was funded by the WM Keck Foundation (MAT and JYY); the office of the OVCR of the University of Hawai’i at Mānoa (JYY), National Institute of General Medical Sciences of the National Institute of Health Grant No. P20GM125508 and P20GM139753 (JYY), Hawaiʻi Community Foundation Grant No. 19CON-95452 (JYY), National Science Foundation Grant No. 2025669 (JYY) and the UHM Undergraduate Research Opportunities Program (AD).

We are grateful to Margaret McFall-Ngai, Edward Ruby, Anthony Amend, Nicole Hynson, Camillo Mora, Craig Nelson, Sean Swift, Pieter Dorrestein, Jannet Janssen, Rob Knight, Eoin Brodie, Jennifer Martiny for helpful scientific discussions; and Tina Carvalho, Kirsten Nakayama, and Nicole Yoneishi for excellent technical assistance. We thank Richard Pezzulo, Chad Durkin, Josie Hoh and Laurent Pool of Hiʻipaka LLC and Waimea Botanical Garden for their generous assistance with providing access to collection sites; and María de la Paz Fernández and Eve Bell for help with the sleep and activity assays and data analysis. All GCMS measurements and HTS library preparations were performed in the Microbial Genomics and Analytical Laboratory core. Experimental fly populations were maintained in the Insectary for Scientific Training and Advances in Research facility. Confocal images were acquired in the UHM Microscopy Imaging Center for Research through Observation.

## Supplemental Data

**Supp. Fig. 1**. Heat map depicting the differential abundance of (**A**) bacterial and (**B**) and fungal genera amongst each population for both early and late generations. Blue labels indicate taxa with significantly different abundances between W1 and W2 populations. Hierarchical clustering is performed with Ward’s method.

**Supp. Fig. 2**. Bacterial (**A-G**) and fungal (**I, J**) taxa found in significantly different abundances between W1 and W2 populations in early (G1-4) and late (G12) generations (Mann-Whitney test). The error bars indicate the maximum and minimum of log-transformed relative abundances; middle line indicates median.

**Supp. Fig 3**. Activity and sleep behavior for W1 and W2 females

**(A)** Daily locomotor activity plots reveal that W1 males are more active during the day compared to W2 males (Mann-Whitney test; n=31 for W1 and W2). The two populations exhibit no differences in nighttime activity. The plots show activity counts in 30 min bins averaged over 3 days and summarized in 24 hrs. The gray shading indicates dark periods.

**(B)** The average activity count for W1 females is higher during the day compared to W2 flies but not at night (Mann-Whitney test). Box plots indicate maximum and minimum values; middle line indicates median. Each point represents the average activity counts per 30 min bin for a single fly over 3 days.

**(C)** The W1 females sleep less during the day compared to W2 flies (Mann-Whitney test).). No differences are found in nighttime sleep levels. The profiles represent average sleep behavior for W1 and W2 females over 3 days, summarized in 24 hrs.

**(D)** The average diurnal sleep duration of W1 flies is significantly shorter than that of W2 females (Mann-Whitney test). Each point represents the sleep duration per 30 min bin for a single fly averaged over 3 days.

**(E)** The W1 and W2 males showed no significant difference in the number of sleep bouts during the day and night (Mann-Whitney test) although W1 males had fewer bouts (mean: 54.7) compared to W2 males (mean: 61.4).

**(F)** The W1 females have fewer diurnal sleep bouts compared to W2 flies (Mann-Whitney test).

**(G)** Axenic W1 and W2 males exhibit similar numbers of sleep bouts during the day and night (Mann-Whitney test).

**(H)** The W1 males have shorter diurnal sleep bouts compared to W2 males (Student’s t-test; values are square root transformed).

**(I)** The W1 females have shorter diurnal sleep bouts compared to W2 females (Student’s t-test; values are square root transformed).

**(J)** Axenic W1 and W2 flies exhibit no detectable differences in sleep bout duration (Students t-test; values are square root transformed).

## Notes

### Competing Interest Statement

The authors have declared no competing interest.

