## Supplementary figures and images for "Environmental microbes promote phenotypic plasticity in *Drosophila* reproduction and sleep behavior"

### Supplemental Figures

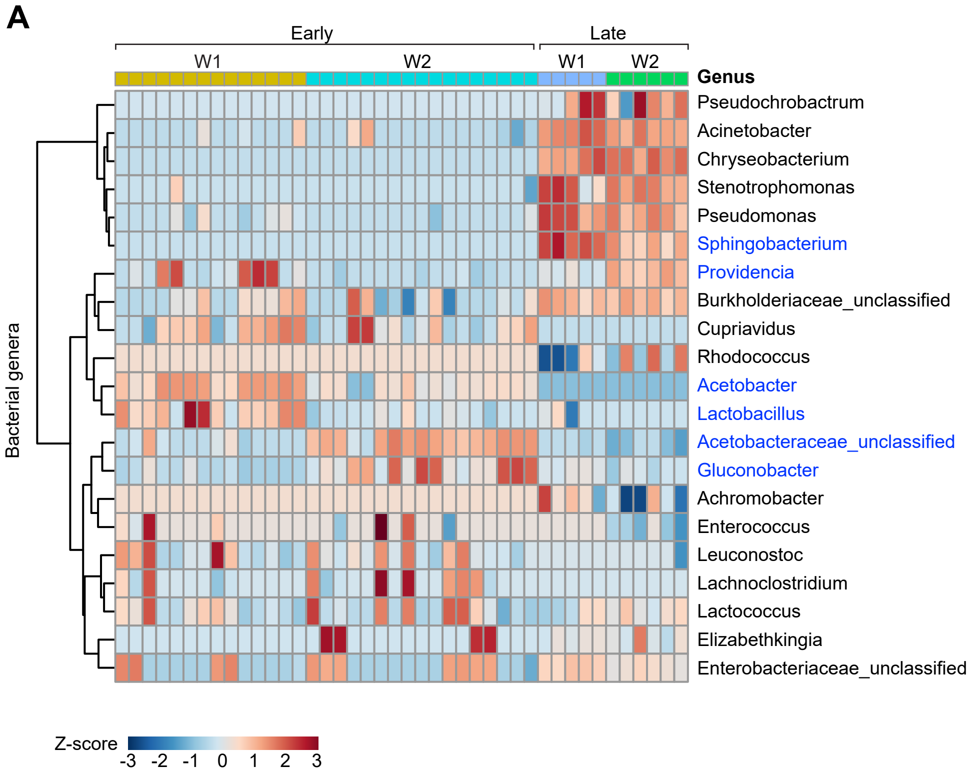

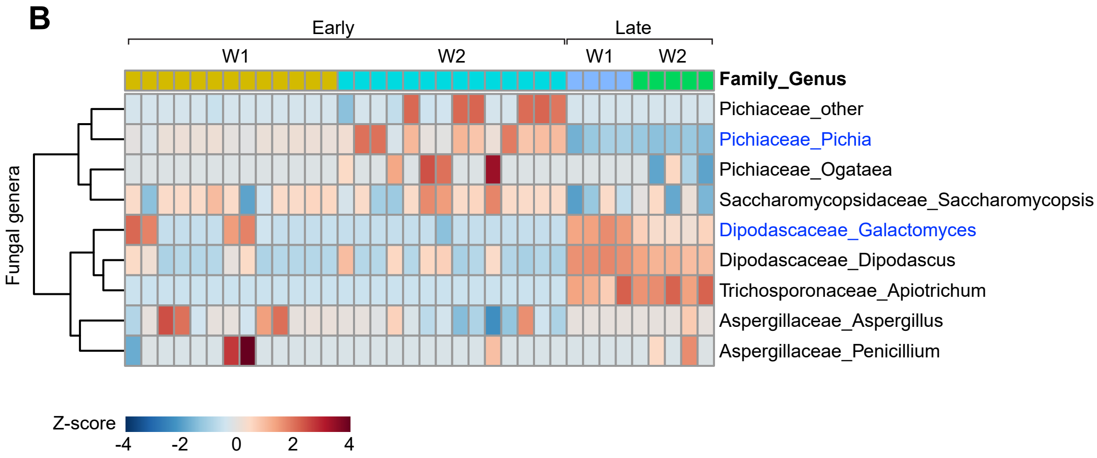

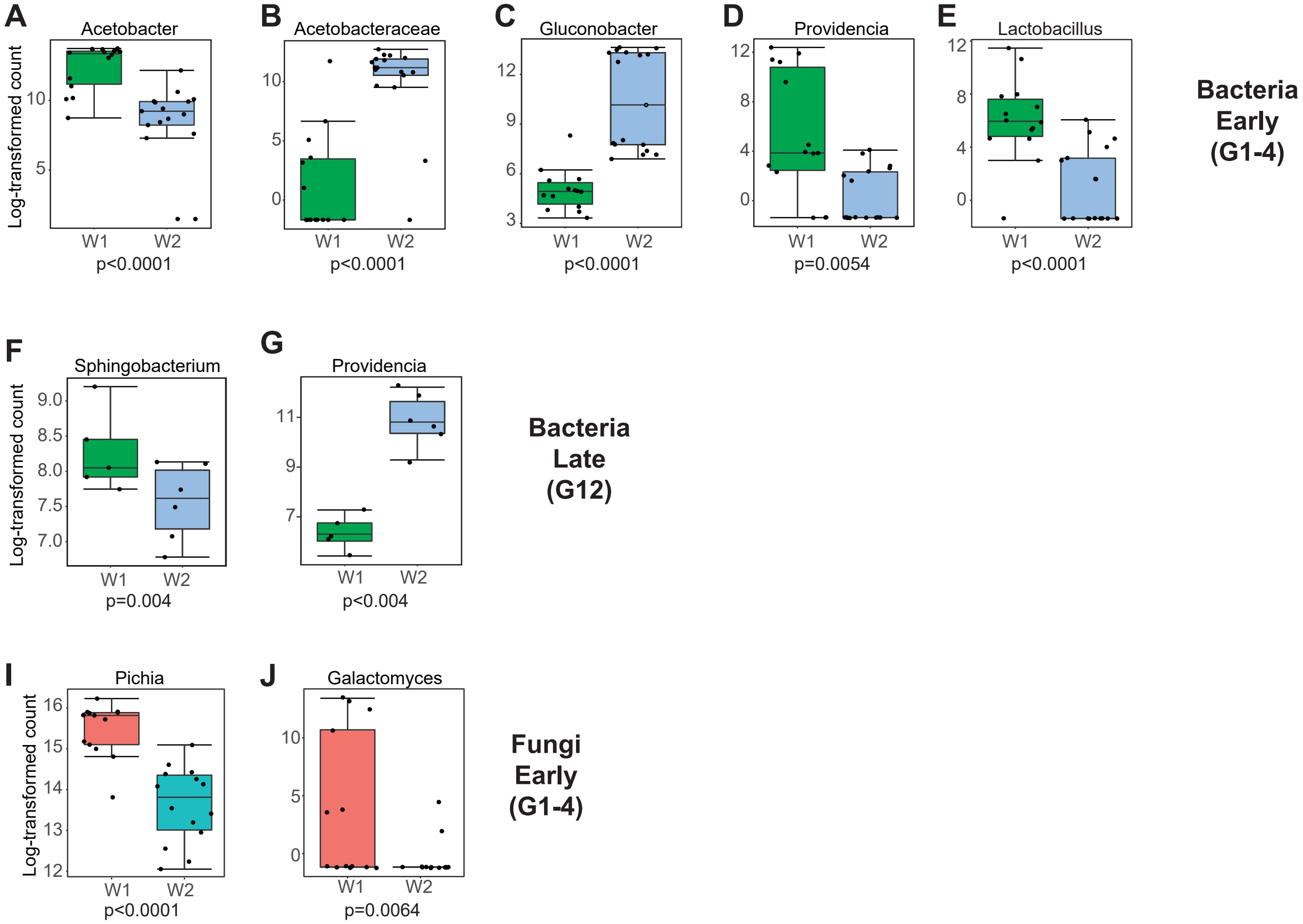

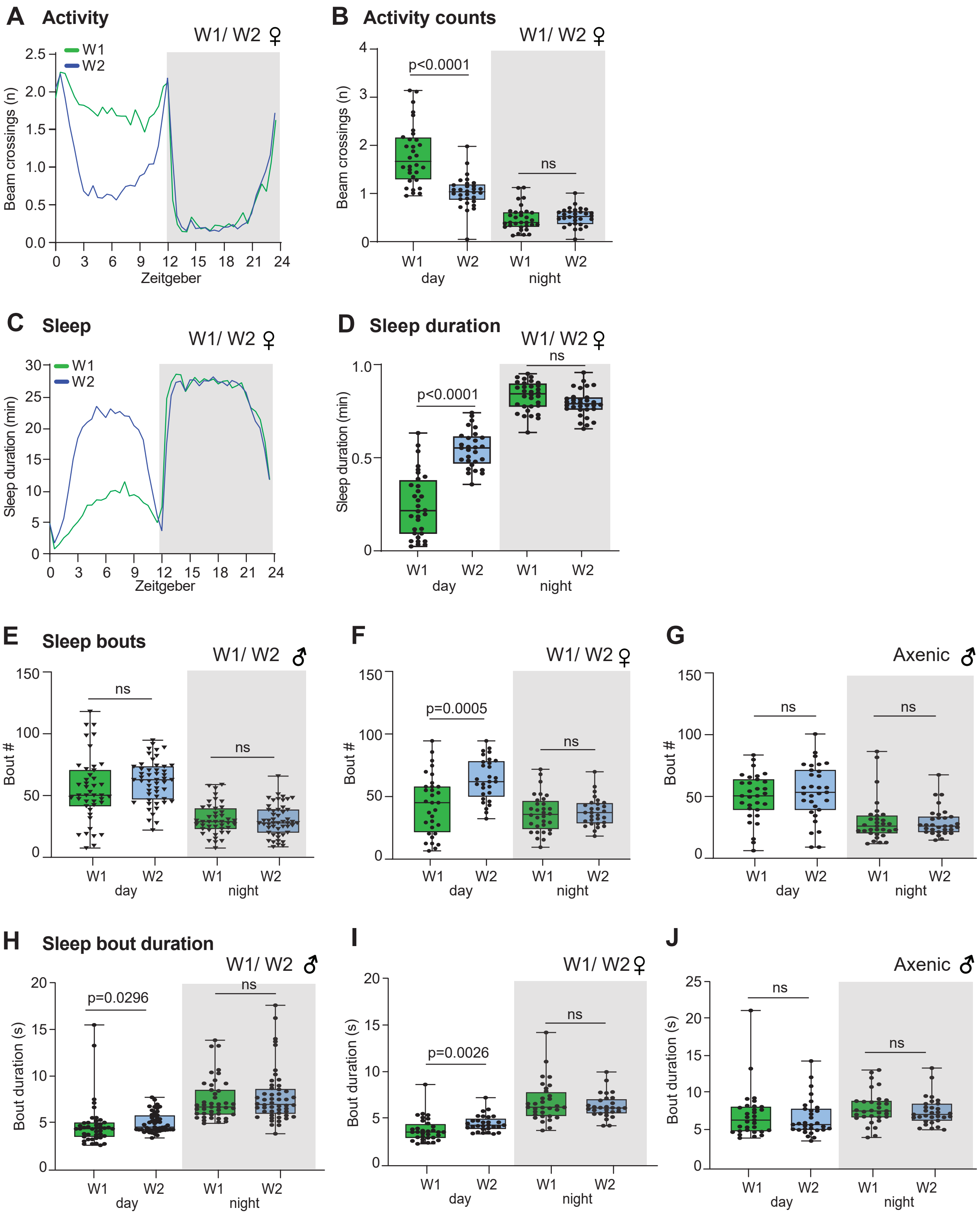
